# IRE-1α is a key switch of pyroptosis and necroptosis in mice by dominating Gasdermin D

**DOI:** 10.1101/2022.10.12.511926

**Authors:** Xin Zeng, Qing Zheng, Deyong Li, Yumeng Li, Qing Xu, Xiaohong Liu, Min Zhu, Wen Liu, Lan Li, Yanrong Lu, Jingqiu Cheng, Younan Chen

## Abstract

Necroptosis and pyroptosis are lytic and inflammatory types of programmed cell death that require the membrane destruction predominantly driven by the mixed lineage kinase domain-like (MLKL) and gasdermin D (GSDMD) proteins, respectively. However, the crosstalk between them remains largely unknown. Here, our research discloses that endoplasmic reticulumn transmembrane protein inositol-requiring enzyme-1α (IRE-1α) is a potential modulator of both necroptosis and pyroptosis, paricularly in liver pathology. Interestingly, enhanced expression of IRE-1α triggers hepatic pyroptosis, while defective IRE-1α level activates hepatic necroptosis, and both processes are closed related to the activity of GSDMD in mice. To elucidate unknown crosstalk, by using pharmacological and genetic methods, we first demonstrated that IRE-1α suppresses necroptosis by promoting the expression of GSDMD and cleaves caspase-8 and by inhibiting the expression of receptor-interacting serine/threonine-protein kinase 1 (RIPK1), RIPK3 and MLKL. Unexpectedly, excess IRE-1α initiates pyroptosis by increasing GSDMD and NLRP3 levels. Our work clearly provides insight into the modulation of IRE-1α to dominate necroptosis and pyroptosis and suggests that IRE-1α may be a promising therapeutic target for drug discovery in both types of tissue injuries.

## Introduction

Pyroptosis is an inflammatory form of programmed cell death and is perhaps most effective in defense against bacteria that invade in hosts without inflammasomes (Shi et al. 2015). Pyroptosis is distinct from other forms of cell death, especially apoptosis, and it drives the activation of caspase-1 or caspase-11/4/5, which cleaves gasdermin D (GSDMD) and release proinflammatory intracellular contents interleukin-1β (IL-1β) and interleukin-18 (IL-18) (Jorgensen and Miao 2015). Importantly, GSDMD is an absolutely necessary effector of pyroptosis, and that co-expresses with canonical inflammasomes invariably includes NOD-like receptor 1 (NLPR1), NLRP3, NLRC4, and AIM2 (Lamkanfi and Dixit 2014). GSDMD separates its N-terminal pore-forming domain (PFD) from the C-terminal repressor domain (RD), and the former oligomerizes to form large pores in the membrane that drive swelling and membrane rupture, which is a critical step mediating pyroptosis (Kovacs and Miao 2017). If the number of GSDMD formed pores in the plasma membrane remains limited normal emergency exocytic membrane fusion events could patch the membrane, similar to MLKL pores from the membrane during necroptosis (Jimenez et al. 2014; Gong et al. 2017). However, the underlying mechanisms by which GSDMD governs pyroptosis are not completely clear, and whether it is involved in the crosstalks of other types of cell death including necroptosis is unknown.

Necroptosis is a highly relevant form of programmed cell death, activated by necrosome, a signaling protein complex that consists of the kinases receptor-interacting serine/threonine-protein kinase 1 (RIPK1) and 3 (RIPK3), and the mixed lineage kinase domain-like (MLKL) (Wang et al. 2014, Rodriguez et al. 2016). It has been discolsed that phosphorylated RIPK3 recruits and activates MLKL, which translocates to the cell membrane and causes membrane rupture (Sun et al. 2012; Zhao et al. 2012). Owing to the loss of membrane integrity in necroptotic cells, the intracellular contents act as damage-associated molecular patterns (DAMPs) that recruit immune cells and cause sterile inflammation in affected tissues (Alvarez-Diaz et al. 2016; Kang et al. 2018). Therefore, necroptosis is considered a highly inflammatory form of cell death, similar to pyroptosis. However, unlike the activation of GSDMD, which is mediated by proteolytic cleavage, a phosphorylation event is essential for MLKL activation during necroptosis. Although both cell death mechanisms invariably rupture the plasma membrane, Annexin V staining of phosphatidylserine, a typical way to detect programmed cell death, is unable to accurately distinguish between pyroptosis and necroptosis, which has been intensively investigated in recent years (Kambara et al. 2018). Moreover, the detailed mechanism that governs necroptotic cell death is largely unknown.

Inositol-requiring enzyme-1α (IRE-1α, also called ERN1) is a sensor of the unfolded protein response (UPR), widely involved in various diseases (Sepulveda et al. 2018). After IRE-1α is activated, the endoribonuclease activity of IRE-1α induces unconventional X-box binding protein 1 (XBP1) splicing. The spliced XBP1 (XBP1-s) controls the upregulation of a broad spectrum of UPR-related genes involved in protein folding, protein entry to the endoplasmic reticulum (ER), and protein quality control (Lee et al. 2002). This process plays an essential role in maintaining ER homeostasis. In addition to its function as a cell fate executor in response to ER stress, IRE-1α is essential in maintaining the intracellular homeostasis of metabolism, reactive oxygen species (ROS), inflammation and cell death (Wang et al. 2012; Ozgur et al. 2015; Duwaerts et al. 2021). It is worth noting that IRE-1α is extremely significant in the liver because hepatocytes are rich in both smooth and rough ER for massive protein synthesis and processing. Although several studies have linked IRE-1α-XBP1 signaling to apoptosis or autophagy (LarabiBarnich and Nguyen 2020; Zhang et al. 2021), the mechanisms connecting its downstream molecules and other types of programmed cell death, such as necroptosis and pyroptosis, remain poorly understood.

In our previous research, we observed the interesting phenomena that either extreme overexpression or suppression of IRE-1α causes liver damage rapidly. Therefore, in this study, we are aiming to further investigate the potential mechanism of IRE-1α and liver cell death using pharmacological and genetic methods. We first demonstrate that IRE-1α is an essential master of pyroptosis and necroptosis by regulating GSDMD. Interestingly, the different expression levels of IRE-1α lead to two distinct effects: overexpression of IRE-1α induces pyroptosis by activating GSDMD and NLRP3, while inhibition of IRE-1α can directly burst necroptosis by reducing GSDMD and cleaved caspase-8, accompanied by intensifying the RIPK3-MLKL pathway. Our findings thus imply that apart from UPR, the equilibrium of IRE-1α is particularly critical for cell survival, since it drives different cell death processes including pyroptosis or necroptosis. Overall, IRE-1α is a potential therapeutic target for treating both pyroptosis or necroptosis associated diseases.

## Results

### Systemic inhibition of IRE-1α leads to acute liver failure

IRE-1α is a major UPR transducer under ER stress, playing a critical role in cell fate determination. To identify the foremost pathway in IRE-1α related cell death, STF083010, a specific chemical IRE-1α inhibitor, was intraperitoneal injected to systemically inhibit IRE-1α in healthy mice (Fig. 1A). To determine the effects of STF083010 (30 mg/kg), the survival rate and the changes in body weight and temperature in each group were monitored throughout the experiment. Surprisingly, the survival rate of the mice markedly decreased shortly after STF083010 treatment, and less than 20% of the mice survived after 72 h (Fig. 1B). Along with the high mortality, a sharp drop in body temperature (< 25°C in the STF083010 group vs. 35°C in the Chow group) was observed beginning 1.5 h after treatment (Fig. 1C). Meanwhile, a substantial decrease in body weight occurred from the first day after STF083010 treatment and worsened day after day, showing a cachectic condition at the end of the experiment (Fig. 1D).

**Figure 1.**
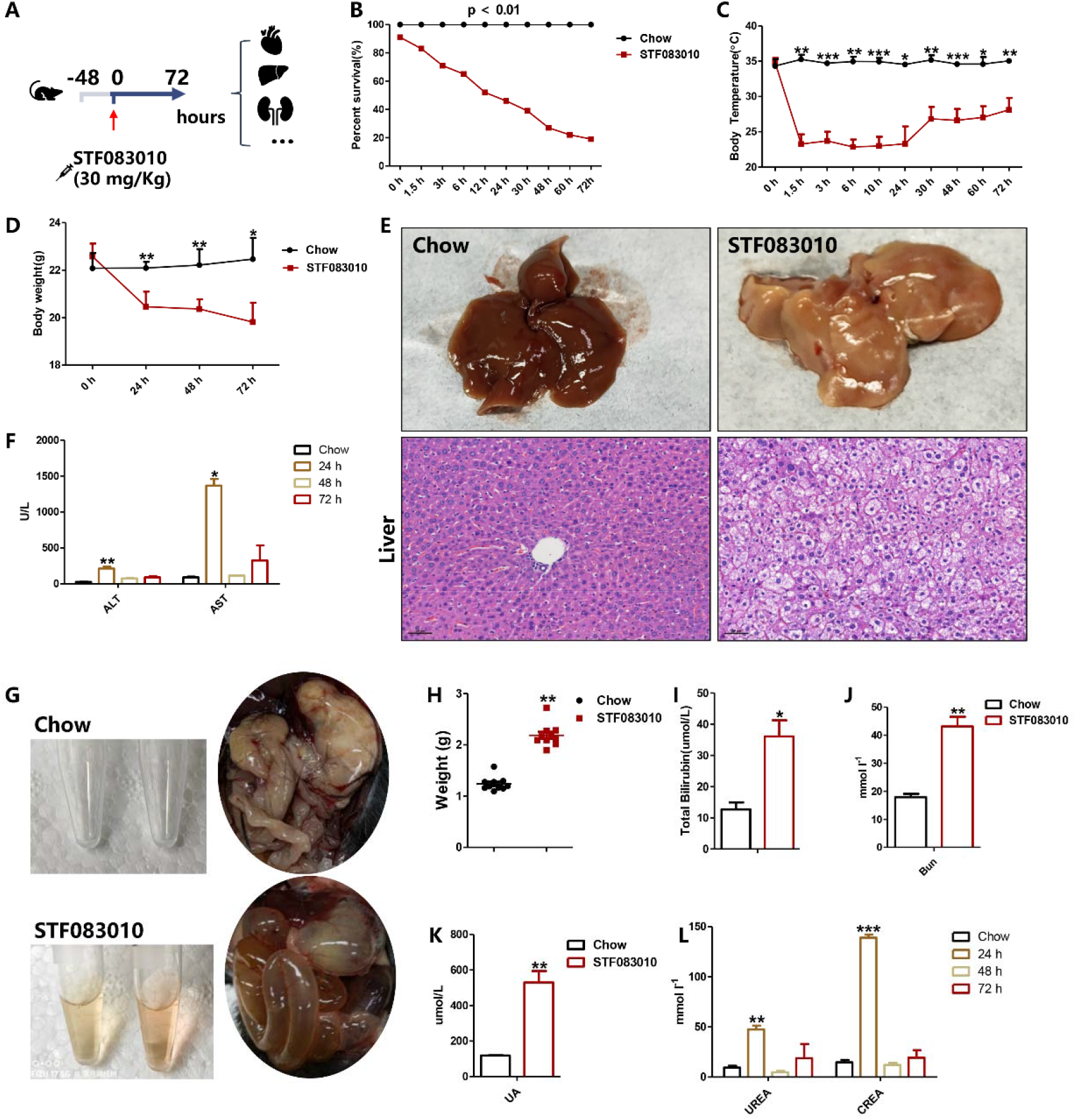
Systemic inhibition of IRE-1α leads to acute liver failure. A Schematic diagram of animal experimental design. STF083010 is a specific chemical inhibitor of IRE-1α. B-D Survival rate, body weight and temperature curves of mice treated with STF083010. n = 40. E Representative images of liver morphology and histology by H&E staining. F Serum ALT and AST levels, n = 10. G Ascites samples of mice and flatulence images of mice treated with STF083010 for 6 h. H-L Biomarkers of acute liver failure in mice, such as liver weight, bilirubin, serum BUN, UA, UREA and CREA levels, n = 10. Data are presented as the means ± SDs. *P < 0.05, **P < 0.01, ***P < 0.001, determined by ANOVA.

To explore the reason of low body temperature in this progression, we observed the pathological changes in liver and skeletal muscle, the main organs of energetic metabolism. After 24 h of STF083010 injection, the livers turned much paler and lacked lustre in comparison with the chow group, and the histology showed cell congestion, edema, degeneration and vacuolization, which indicated acute massive hepatocyte injury (Fig. 1E). In addition, severe ascites and flatulence, as well as liver weight increase were found (Fig. 1G and H), and blood biochemistry indicated that the plasma levels of ALT, AST and total bilirubin were significantly elevated, indicating serious abnormalities in liver function in the STF083010 group (Fig. 1F and I). Together with the indications of acute liver failure (ALF), serum UREA, CREA, and BUN were distinctly increased, suggesting that severe hepatorenal syndrome occurred after systemic IRE-1α inhibition (Fig. 1G - L). In contrast to the severe pathological changes in the liver, we checked the morphology and histology of skeletal muscle, but no observable abnormalities were found (Fig. S1a and b). Similarly, no obvious histological change was found in other tissues, including the heart, lung and colon (Fig. S1c - h).

Moreover, to confirm the detrimental effects of IRE-1α deficiency on cells, mouse hepatocyte and myoblast cell lines (AML-12 and C2C12, respectively) were exposed to different concentrations of STF083010 for 24 h. Cell viability was measured by CCK8 assay and the results suggested that the viability of AML-12 cells was markedly deteriorated in a dose-dependent manner, but C2C12 cells showed no obvious change (Fig. S1i and j). In addition, STF083010 did not damage cell viability of other cell lines such as renal tubular cell line HK-2 and endothelial cells HUVECs (Fig. S1k - p). In summary, these results demonstrated that inhibition of IRE-1α was able to powerfully damage hepatocytes and induce severe acute liver injury, leading to high mortality in mice.

### Necroptosis is possibly the main cause of hepatic cell death induced by STF083010

To explore the mechanisms underlying IRE-1α deficiency-induced liver injury, we focused on the identification of the cell death form in the liver after STF083010 treatment. First, the TUNEL results of the mouse liver samples showed that inhibition of IRE-1α increased cell death in the STF083010 group at 24 h (Fig. 2A). Since the TUNEL assay is unable to distinguish different programmed cell death processes, among apoptosis, necroptosis and pyroptosis, we examined specific molecular markers of these cell death forms. The qPCR results of liver samples clearly showed that STF083010 did not alter the mRNA expressions of apoptotic genes *Caspase-3, Caspase-6* and *Caspase-9*, the most important terminal cleavage enzymes in apoptosis (Fig. 2B). Meanwhile, activated caspase-3/-6/-9 were barely altered, as shown by Western blotting (Fig. 2E). However, STF083010 significantly upregulated the mRNA expression of *Nlrp3*, a typical marker of inflammasomes, after 24 h of treatment, but downregulated *Gsdmd*, an important effector of pyroptosis (Fig. 2C). Consistently, the Western blot indicated a distinct elevation of NLRP3 but a decrease in active GSDMD level, while the protein level of caspase-1 did not show significant alteration. This phenomenon is unexpectedly different from previous studies in which simultaneous increases in NLRP3 and GSDMD were reported in patients or animal models of NASH or diabetes mellitus (Fig. 2F). More importantly, in addition to the changes in pyroptosis markers, we found that the mRNA expressions of necroptosis markers, including *Ripk1, Ripk3* and *Mlkl*, were prominently increased (Fig. 2D). Consistently, the STF083010 group displayed decreased protein levels of cleaved caspase-8 and the most distinct upregulation of p-MLKL and p-RIPK3 (Fig. 2G).

**Figure 2.**
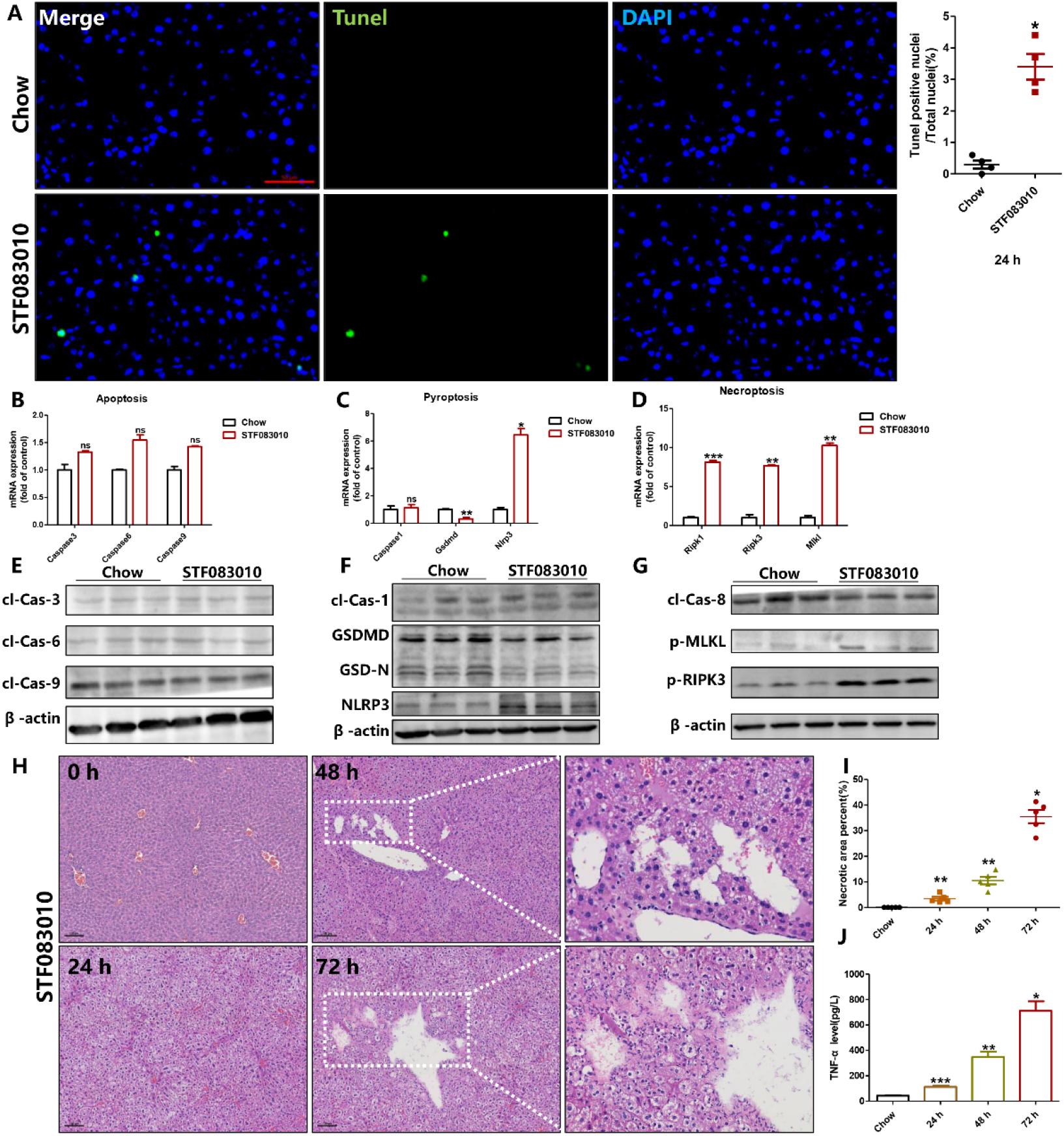
Necroptosis is possibly the main cause of hepatic cell death. A Staining for TUNEL (green) and DAPI (blue) (scale bars; 50 μm). n = 3. B-D The mRNA expression levels of apoptotic genes, including *Caspase-3, Caspase-6* and *Caspase-9*; pyroptotic genes, including *Caspase-1, Gsdmd, Nlrp3*; and necroptosis markers, including *Ripk1, Ripk3* and *Mlkl*, were detected after STF083010 treatment for 24 h, and β-actin was used as an internal control. n = 5. E-G Representative western blots of apoptotic markers, including cleaved Caspase-3/-6/-9; pyroptotic markers, including Caspase-1, GSDMD and NLRP3; and necroptosis markers, including cleaved Caspase-8, p-RIPK3 and p-MLKL, were detected after 24 h of treatment, and β-actin was used as a protein loading control. n = 3. H-I Representative images of H&E-stained liver tissues after 24-72 h of treatment with STF083010. Dotted lines in H&E images outline necrotic areas. n = 5. J The levels of TNF-α in serum. n = 5. Data information: Data are presented as the means ± SDs. *P < 0.05, **P < 0.01, ***P < 0.001, determined by ANOVA.

With respect to the long-term consequences, the liver histology of the STF083010 group showed massive hepatocyte injury and spotty necroptosis at 48 h and 72 h. ALF mice showed areas of confluent hemorrhagic necrosis, congestion and cell perforation-induced autolysis (Fig. 2H and I, dotted line). Furthermore, the release of TNF-α in serum was robustly increased in line with the exposure time and peaked at 72 h (Fig. 2J), which confirmed the activation of necroptosis. In summary, our results showed that necroptosis may be the dominant form of hepatic cell death after IRE-1α inhibition and that NLRP3 or GSDMD may be related to the process.

### IRE-1α dominates the switch between pyroptosis and necroptosis in hepatocytes

To further elucidate the link between IRE-1α and cell death in hepatocytes, human hepatocyte cell line LO-2 cells and mouse primary hepatocytes (MPH) were exposed to different concentrations of STF083010 for 24 h. Consistent with the in vivo results, STF083010 markedly deteriorated cell viability in a dose-dependent manner (Fig. S2a and b). Next, we examined specific molecular markers of necroptosis and pyroptosis. The qPCR results clearly showed that *Gsdmd* was downregulated, but *Nlrp3, Ripk1, Ripk3* and *Mlkl* were significantly upregulated in LO-2 cells (Fig. 3A and B) and MPH cells (Fig. 3C and D). Additionally, the protein expression levels corresponded with the mRNA expression panel and also agreed with the relevant results in mouse models, showing no obvious change in apoptotic markers, but significantly increased levels of NLRP3, p-MLKL and p-RIPK3, and decreased levels of GSDMD and cleaved caspase-8 in both LO-2 cells (Fig. 3E) and MPH cells (Fig. 3F) after STF083010 treatment. Meanwhile, to convince the effects of STF083010, we used different specific inhibitors of IRE-1α, such as 4u8c and APY29, to treat LO-2 cells. The results were in consistent with STF083010, revealing that inhibition of IRE-1α significantly upregulated the mRNA expression of the necroptosis markers *Ripk1* and *Mlkl*, but downregulated *Gsdmd* (Fig. 3G). However, the combination of STF083010 with low-dose LPS treatment, a common pyroptosis inducer that elevates GSDMD levels, was able to robustly reverse these changes, leading to induction of GSDMD but suppression of p-MLKL remarkably, as evidenced by an in situ immunofluorecence assay with GSDMD or p-MLKL antibodies (Fig. 3I and J).

**Figure 3.**
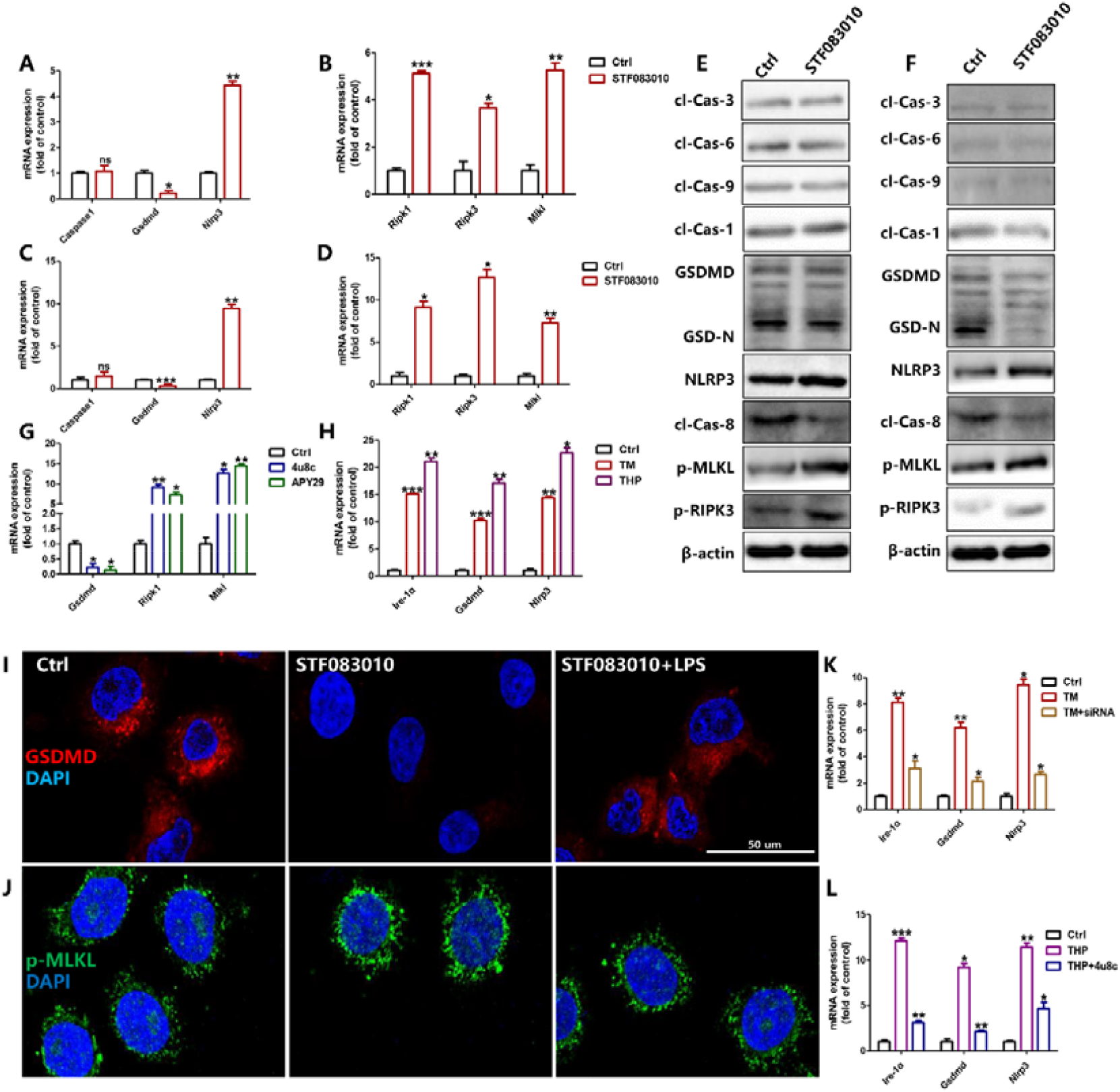
IRE-1α dominates the switch between pyroptosis and necroptosis in hepatocytes. A-D The mRNA expression of pyroptotic and necroptotic genes was detected after STF083010 treatment for 24 h, and β-actin was used as an internal control. n = 3. E, F Representative western blots of apoptotic, pyroptotic and necroptosis markers after 24 h of treatment, and β-actin was used as a protein-loading control. n = 3. G, H The mRNA expression of *Ire-1α, Gsdmd* and *Nlrp3* was detected after TM or THP and 4u8c or APY29 treatment for 24 h, and β-actin was used as an internal control. n = 3. I, J Cells were exposed to STF083010 and labeled by immunofluorescence with anti-GSDMD (red), p-MLKL (green), and DAPI (blue) (scale bars; 50 μm), respectively. n = 3. K, L The mRNA expression of *Ire-1α, Gsdmd* and *Nlrp3* was detected after TM and siRNA or THP and 4u8c treatment for 24 h, and β-actin was used as an internal control. n = 3. Data information: Data are presented as the means ± SDs. *P < 0.05, **P < 0.01, ***P < 0.001, determined by ANOVA.

A great number of studies have demonstrated that IRE-1α can be chemically induced by some ER stressors, such as tunicamycin (TM) and thapsigargin (THP). TM induces ER stress by inhibiting the glycosylation of newly synthesized proteins, and THP induces ER stress by inhibiting Ca^2+^-ATPase, which destroys Ca^2+^ homeostasis in the ER. In our previous experiments, we have demonstrated that inhibiting IRE-1α induced necroptosis, along with downregulated GSDMD. In order to further elucidate the role of IRE-1α in programmed cell death, as well as the relationship between IRE-1α and GSDMD, LO-2 cells were exposed to TM or THP for 24 h. As expected, we found that ER stressors increased the expression of *Ire-1α*, and consequently increased the expressions of pyroptosis markers *Gsdmd* and *Nlrp3* (Fig. 3H). Besides, siRNA of IRE-1α was used in MPHs cells challenged with of TM for 24 h. The qPCR results clearly showed that knockdown of IRE-1α remarkably repressed the pyroptotic markers GSDMD and NLRP3 (Fig. 3K), and the same results appeared when MPHs were treated with THP with or without 4u8c (Fig. 3L). These data displayed that increased IRE-1α might activate pyroptosis, but inhibition of IRE-1α apparently reverse this process.

Given that, we hypothesized that IRE-1α is capable of regulating GSDMD, which may govern the cell fate, triggering the switch between necroptosis and pyroptosis. To visualize the cell death process, AO/EB staining was utilized to specifically observe the changes in cell permeability under a microscope (Fig. 4A).

**Figure 4.**
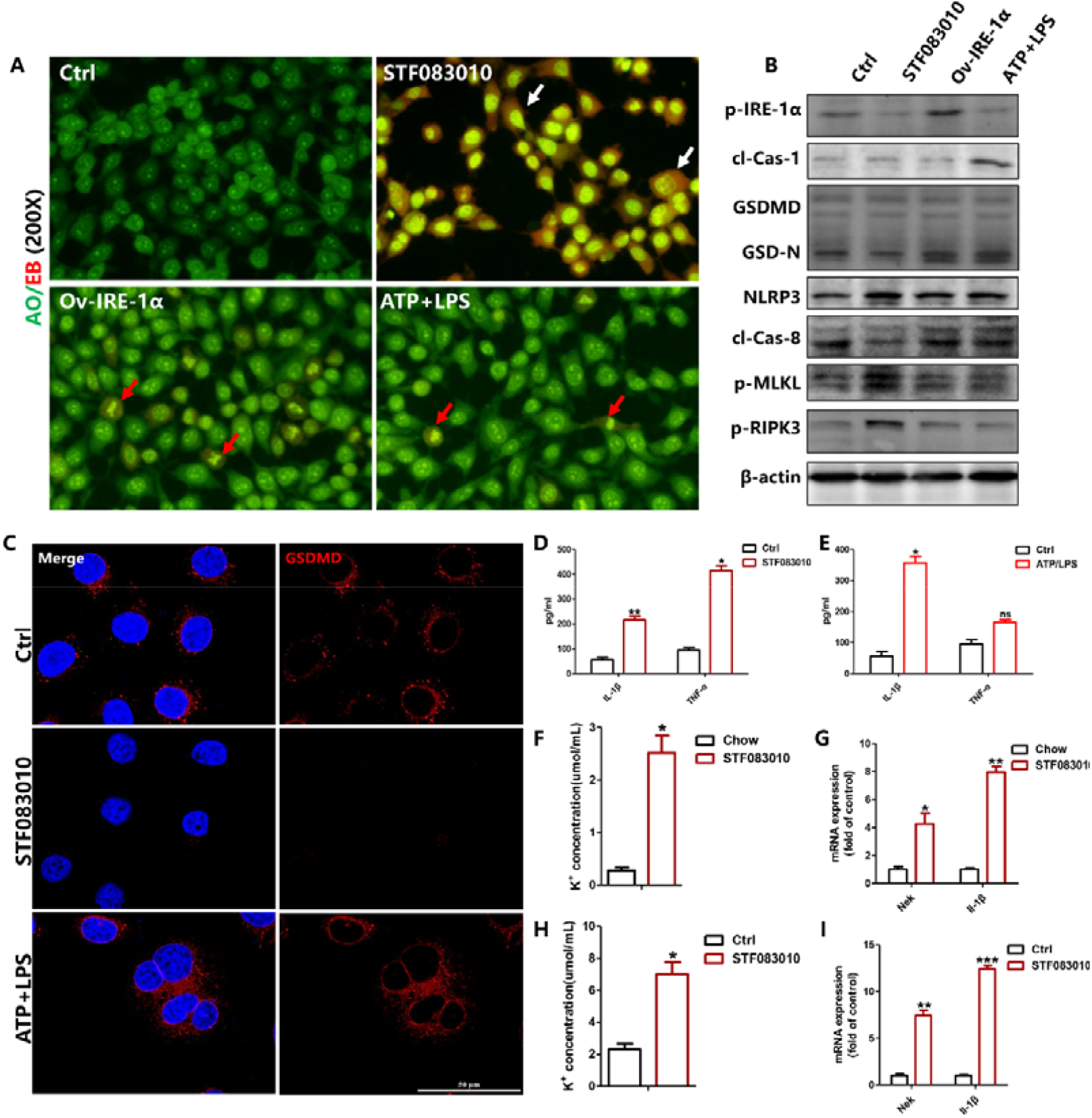
GSDMD is a negative executor of necroptosis. A The morphology of cells was visualized by AO/EB staining. All groups were visualized under a microscope at 200× magnification after 24 h of treatment. n = 3. B Representative western blots of key proteins after 24 h of treatment, and β-actin was used as a protein-loading control. n = 3. C Cells were exposed to STF083010 or ATP/LPS and subsequently labeled by immunofluorescence with anti-GSDMD (red) and DAPI (blue) (scale bars; 50 μm), respectively. n = 3. D, E The levels of IL-1β and TNF-α in the supernatant of STF083010- or ATP/LPS-treated LO-2 cells. n = 3. F, H Potassium concentration in serum and in the supernatant after STF083010 treatment. n = 8. G, I The mRNA expression of mice after STF083010 treatment for 24 h. n = 8. Data information: Data are presented as the means ± SDs. *P < 0.05, **P < 0.01, ***P < 0.001, determined by ANOVA.

The results suggested that STF083010-treated cells were obviously swollen and had red fluorescence overload in the cytoplasm and bright yellow in the enlarged nuclear (white arrow), compared with cells of other groups, which suggested that more EB entered into the cells, probably by the p-MLKL pore. More importantly, a plasmid was used to overexpress IRE-1α in LO-2 cells, and the morphological alteration of plasmid-treated cells was more similar to pyroptotic cells after ATP/LPS exposure that with little red fluorescence and nuclear fragmentation (red arrow) (Fig. 4A). Consistently, the Western blot results indicated that STF083010 treatments substantially increased the protein levels of the necroptosis markers p-MLKL and p-RIPK3, but overexpression of IRE-1α obviously upregulated the pyroptotic markers GSDMD and NLRP3 (Fig. 4B). These results suggested that IRE-1α may dominate the switch between pyroptosis and necroptosis. Overexpression of IRE-1α elicits pyroptosis, while deficiency of IRE-1α switches cells fate to necroptosis.

### GSDMD maybe a negative executor of necroptosis

GSDMD was discovered to be able to form pores on the cell membrane, acting as an effector for pyroptosis. Meanwhile, NLRP3 inflammasome activates procaspase-1 to be cleaved into p20 and p10 subunits that form the active caspase-1, which then leads to maturation and secretion of IL-1β and IL-18 in pyroptosis. Nevertheless, the roles of GSDMD and NLRP3 in necroptotic cell death are largely unknown. To visualize this alteration intuitively, we used immunofluorescence to confirm that STF083010 profoundly prevented the expression of GSDMD in LO-2 cells, meanwhile, as a positive control, the typical pyroptosis inducer ATP and LPS markedly increased the expression of GSDMD in LO-2 cells (Fig. 4C and Fig. S2c). Another study found that STF083010 induced more release of TNF-α than IL-1β, indicating necroptosis was activated (Fig. 4D), but conversely, ATP and LPS stimulated more IL-1β than TNF-α, indicating pyroptosis was activated (Fig. 4E).

Many studies have shown that NLRP3 senses a variety of stimuli, such as toxins, pathogens, metabolites, crystalline substances, nucleic acids, and ATP, and always shows coordinated alterations with GSDMD in various inflammatory diseases clearly. Moreover, it was previously demonstrated that membrane-associated MLKL-induced potassium efflux is able to induce NLRP3 signaling in necroptosis. Therefore, the potassium concentration in serum or in the cell culture supernatant was detected, and the results showed that the content of K^+^ increased after STF083010 treatment (Fig. 4F and G). Furthermore, the mRNA expression of *Nek*, as an upstream trigger of NLRP3 inflammasome formation, and *Il-1β*, as a marker of the activated NLRP3 inflammasome, were significantly upregulated (Fig. 4H and I), which paralleled the upregulation of NLRP3 upon necroptosis (Fig. S2d). These results suggested that enhanced potassium efflux in necroptosis may induce the expression of NLRP3, so we speculated that increased NLRP3 may be the outcome rather than driver of necroptosis underlying IRE-1α inhibition. Consequently, we further hypothesized that IRE-1α may dominate the switch between pyroptosis and necroptosis by governing the activity of GSDMD, that may play a significant role and probably be a negative executor of necroptosis.

### IRE-1α equilibrium improves mice survival

Next, we further explored whether IRE-1α expression governs necroptosis or pyroptosis in vivo. In our previous study, TM was found able to induce the overexpression of IRE-1α, and activate ER stress and inflammation, resulting steatosis in animal models. Thus, in this study, a lower dose (10 mg/kg) of STF083010 was used three days before TM (2 mg/kg) treatment in mice to inhibit the overexpression of IRE-1α-induced pyroptosis (Fig. 5A). The results showed that the plasma levels of ALT, AST, CREA and UREA were strikingly increased in the TM group, suggesting that TM-induced ER stress drove acute liver damage; however, pretreatment with a low dose of STF083010 before TM administration effectively decreased the above biochemical markers and substantially ameliorated liver function (Fig. 5B and C). Furthermore, the livers rapidly recovered from TM-induced steatosis, as evidenced by liver morphology and Oil Red O staining (Fig. 5D). Similarly, the release of IL-1β and IL-18 in serum indicated that pyroptosis was prominently activated after TM challenge; nevertheless, STF083010 reversed this activation effectively (Fig. 5E and F). Consistently, pyroptotic markers such as IRE-1α and GSDMD were enhanced in the liver after TM treatment, but these molecules were effectively downregulated in the low-dose STF083010 pretreatment group. However, the key factors of necroptosis decreased slightly in the TM group, and there was a slight increase after pretreatment with a low dose of STF083010 (Fig. S3a, b). Overall, these results revealed that the excessive IRE-1α unambiguous boosted pyroptosis in the liver attributable to profound ER stress, but maintaining the IRE-1α balance by proper inhibition of IRE-1α by using a low dose of STF083010 pretreatment effectively improved mouse survival.

**Figure 5.**
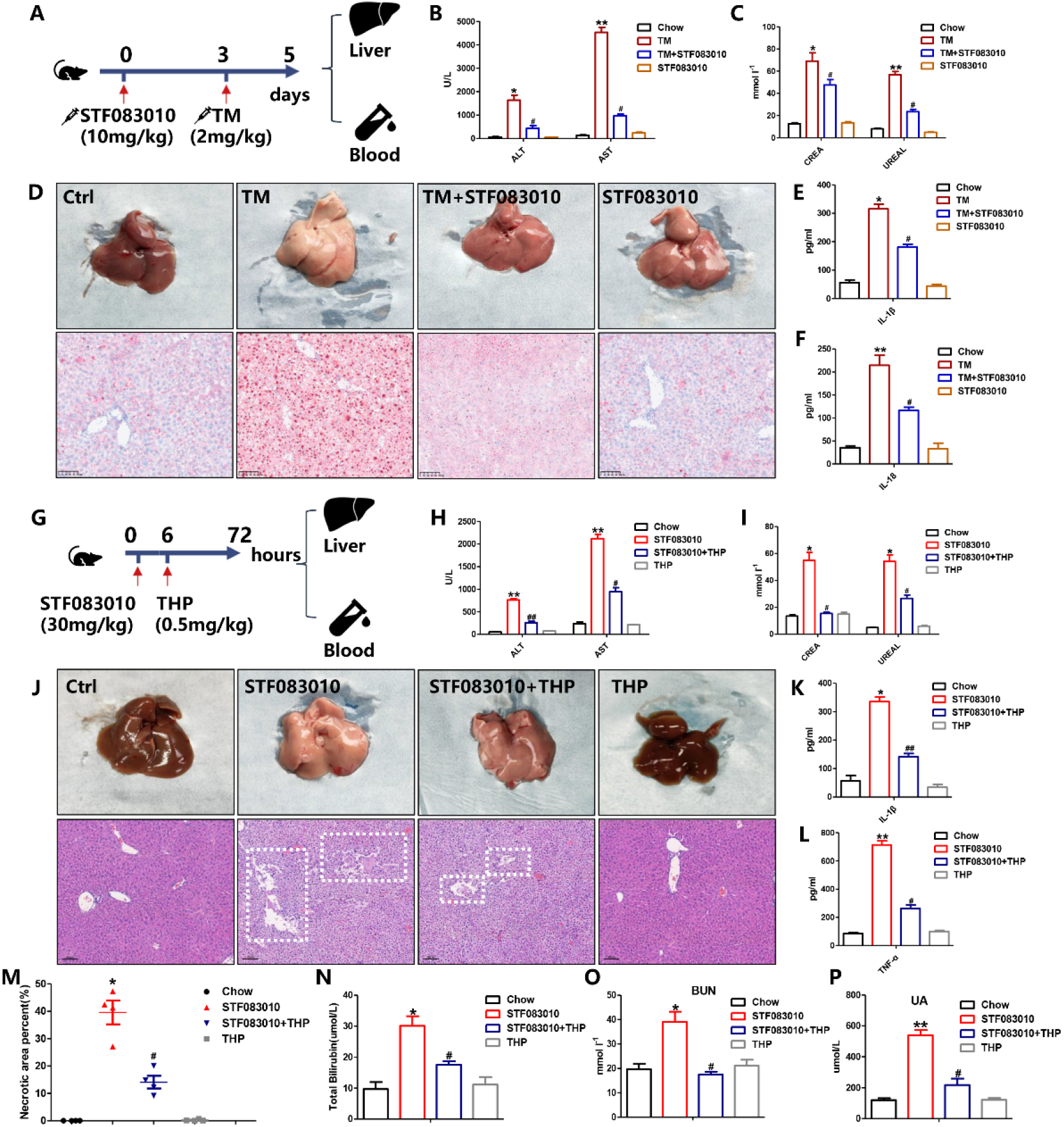
The IRE-1α equilibrium improves mouse survival. A Schematic showing that mice were treated with TM and/or STF083010. n = 10. B, C Serum ALT, AST, UREA and CREA levels. n = 8. D Representative liver and Oil Red O-staining. n = 8. E, F the levels of IL-1β and TNF-α in serum of TM and/or STF083010–treated mice. n = 5. G Schematic showing that mice were treated with STF083010 and/or THP. n = 10. H, I Serum ALT, AST, UREA and CREA levels. n = 8. J Representative liver and H&E-staining. Dotted lines in H&E images outline necrotic areas. n = 6. K, L the levels of IL-1β and TNF-α in serum of STF083010 and/or THP–treated mice. n = 5. M-P Biomarkers of acute liver failure, such as necrotic areas and bilirubin, serum UA and BUN levels. n = 8. Data information: Data are presented as means ± SDs. *P and ^**#**^P< 0.05, **P and ^**##**^P< 0.01, determined by ANOVA.

Meanwhile, to further confirm whether defective IRE-1α is devoted to necroptosis, we used high-dose STF083010 (30 mg/kg) to suppressIRE-1α, and induce acute liver failure and necroptosis, then used THP to restore the IRE-1α level (Fig. 5G).

Consistent with our previous results, the plasma levels of ALT, AST, CREA and UREA were distinctly increased in the STF083010 group; nevertheless, low-dose THP (0.5 mg/kg) alone did not show obvious detrimental effects on liver function, but when combined with STF083010, it significantly improved hepatorenal syndrome (Fig. 5H and I). Moreover, liver histology showed that overexpression of IRE-1α was enough to improve necroptosis-induced massive hepatocyte injury and spotty areas (Fig. 5J and M). Consistently, the related biomarkers, such as bilirubin, BUN and UA, decreased after THP exposure (Fig. 5N, O and P). In addition, the markedly increased serum levels of IL-1β and TNF-α were substantially suppressed by THP treatment (Fig. 5N, K and L). Similarly, necroptotic markers such as RIPK1, RIPK3 and MLKL in the liver were significantly increased after STF083010 treatment, but the key factors of pyroptosis were markedly decreased. Nevertheless, IRE-1α, GSDMD and cleaved Caspase-8 were obviously upregulated in the STF083010 + THP group, and necroptotic markers were visually decreased (Fig. S3c, d). These data suggested that the low-dose THP-induced UPR may effectively increase IRE-1α expression to some extent, which may compensate for high-dose STF083010-driven IRE-1α-GSDMD deficiency and reverse severe hepatic necroptosis-induced liver injuries. In summary, excess or deficient IRE-1α determines the alternative form of cell death, and maintaining the balance of IRE-1α is important for liver function and mouse survival.

## Discussion

IRE-1α initiates the most conserved signaling branch of the UPR, and its role in disease has been extensively discussed (Sepulveda et al. 2018). However, the downstream signalings of IRE-1α are poorly understood. Previous studies have disclosed the function of IRE-1α, and found that the IRE-1α-XBP1 pathway is invariably a primary consideration. Many studies have demonstrated that IRE-1α splices XBP1 to produce a potent transcription factor (XBP1s) that regulates the expression of genes important for ER homeostasis (Hetz and Glimcher 2009). In this context, several studies have further explored that IRE-1α was proposed to act through the activation of JNK, whereby it controls the activities of apoptosis and autophagy under ER stress (Kanda and Miura 2004). Similarly, IRE-1α signaling has recently been linked to the expression of proapoptotic BCL-2 family members and ASK1-interacting protein 1 (AIP1) (Kim et al. 2009). Despite all this, IRE-1α is little known in other cell death pathways, such as pyroptosis and necroptosis, even though it has been extensively studied in apoptosis and autophagy. Here, we have foremost uncovered GSDMD as another candidate target of IRE-1α that clearly differs from XBP1, and consequently the cleavage of GSDMD by inflammatory caspases sorely determines pyroptosis. Here we demonstrated that the overexpression of IRE-1α may increase the susceptibility of cells to pyroptosis by upregulating GSDMD, however, unexpectedly, inhibition of IRE-1α may synergistically suppress GSDMD, leading to serious activation of necroptosis, suggesting IRE-1α-GSDMD axis plays a central role in cell fate determination, particularly in liver pathology. This conclusion was fully confirmed in both mouse models and in vitro cell experiments, which enlarges our knowledge of IRE-1α for future research.

GSDMD is one of six gasdermins, which are a conserved family of proteins expressed in various tissues and mainly function to regulate cell death. It is well demonstrated that GSDMD-pore formation mediates pyroptosis (Kovacs and Miao 2017). Recent studies have revealed that GSDMD is cleaved by caspase-4/5/11 activation or Pannexin-1 deficiency, resulting in the efflux of K^+^ and inducing the assembly of the NLRP3 inflammasome, eventually leading to pyroptosis in the noncanonical pathway (HoffmannLambert and Pedersen 2009). More importantly, data have shown caspase-8-mediated cleavage of GSDMD also leads to pyroptosis in the same pathway, and other study showed that cleavage activity of caspase-8 is in turn modulated by GSDMD (Sarhan et al. 2018; Chen et al. 2019). Therefore, it is also possible that downregulation of GSDMD can affect caspase-8 activity and remodel the cell death pathway. Although it has also been demonstrated that gasdermins can convert apoptosis into pyroptosis and vice versa, we showed another possibility that GSDMD suppression may elicit necroptosis by raising RIPK1, RIPK3 and MLKL. The detailed mechanisms need to be explored further. Moreover, it is understandable that p-MLKL pore-mediated K^+^ efflux leads to NEK7 binding to NLRP3 to trigger its oligomerization, which is a reasonable explanation of the high NLRP3 level found in low GSDMD-induced necroptosis. In addition, though GSDMD is the first effector protein to be identified in pyroptosis and has been studied the most frequently, the perspicuous mechanism of its roles in cell death is far from full known. IFN regulatory transcription factor-2 (IRF2) has been reported to transcriptionally regulate GSDMD expression (Liao et al. 2019). Therefore, our study suggested IRE-1α is one of the most powerful modulators of GSDMD.

Pyroptosis was first discovered in 1992 and was defined as proinflammatory programmed cell death in 2001, which distinguishes it from apoptosis (Zychlinsky and Sansonetti 1992; Cookson and Brennan 2001). Pyroptosis is accompanied by inflammasome activation, leading to cytokine release in multiple tissues and impacting a variety of disorders, such as tumors, diabetes, and obesity (Du T et al. 2021). Pyroptosis is a double-edged sword and plays an important role in tumor development. Previous work found that GSDMA was a tumor suppressor gene in gastric cancer (Saeki et al. 2009), but GSDMB was overexpressed in some gastric cancer cells and could act as an oncogene (Komiyama et al. 2010), which was similar to breast cancers (Hergueta-Redondo et al. 2014). The increase in GSDMC in colorectal cancer cells (Miguchi et al. 2016) and GSDMD were found to be upregulated in non-small cell lung cancer (Gao et al. 2018), but GSDME generally exerted a tumor suppressive effect (de BeeckVan Laer and Van Camp 2012). Various metabolic danger signals activate pyroptosis, such as uric acid crystals in gout, cholesterol crystals and oxidized low-density lipoprotein (ox-LDL) in atherogenesis, and glucose, fatty acids and islet amyloid polypeptide in type 2 diabetes (Renaudin et al. 2020; He et al. 2021; Zhou et al. 2010). More importantly, our previous results revealed that olive oil treatment effectively abolished saturated fatty acids induced deleterious effects in long term high-fat diet induced NAFLD by alleviating pyroptosis in liver (Zeng et al. 2020). In this study, the data revealed that IRE-1α might trigger GSDMD- and NLRP3 inflammasome-mediated pyroptosis activation. Consequently, the relationship between IRE-1α and metabolic disorders, such as NAFLD and diabetes, has become increasingly understood. The enlarged knowledge provides some inspirations for further clinical treatments towards these diseases.

Necroptosis is a form of regulated necrotic cell death mediated by RIPK1, RIPK3 and MLKL (Rodriguez et al. 2016), while their regulatory signals are rarely reported to date. Several studies have demonstrated that necroptosis is activated by TNF-α signaling, one of the mostly studied proinflammatory cytokines. Significantly, necroptosis causes proinflammatory responses, releasing cytokines, including TNF-α and IL-1β, which have been proven to be key cytokines in the development of coronavirus disease 2019 (COVID-19) (Zhang et al. 2020). It is not surprising that necroptosis is involved in not only inflammatory diseases, but also various other diseases (Shan et al. 2018). For example, in ischemia/reperfusion (I/R), Ripk3^-/-^ mice are less sensitive to heart failure, while signs of an ongoing necroptotic response are common to various renal and liver disorders (Linkermann et al. 2012; Cao and Mu 2021). In our study, we found the inhibition of IRE-1α-GSDMD-caspase 8 pathway contributes to the activation of necroptosis in liver, and the agreeable evidence exists that ablation of caspase 8 strongly aggravated liver injury and fibrosis in the methionine–choline-deficient (MCD) diet-induced model, and simultaneously RIPK3 was strongly upregulated in this model. More importantly, patients with NASH had a dramatic upregulation of hepatic RIPK3 expression as an indicator of necroptosis (Gautheron et al. 2014; Zhang et al. 2019). Apart from above findings, the role of necroptosis in hepatocarcinogenesis is less clear. A latest report showed unexpected results that pure necroptosis activation suppresses inflammation, proliferation and carcinogenesis in the liver, and the suppression of necroptosis reverts the necroptosis-dependent cytokine microenvironment and switches intrahepatic cholangiocarcinoma to hepatocellular carcinoma rapidly (Seehawer et al. 2018). Taken together, here we found that IRE-1α inhibits necroptosis probably through GSDMD and caspase 8 signaling pathway, but the underlying regulatory mechanisms need to be further explored.

In summary, our study identified IRE-1α as an important regulator governing programmed cell death in liver. The equilibrium of IRE-1α coordinates with GSDMD activity, determining the switch between pyroptosis and necroptosis. The overexpression of IRE-1α may increase the susceptibility of cells to pyroptosis, by increasing GSDMD and inflammasome activities; while IRE-1α deficiency may trigger liver necroptosis, which is possibly attributable to the inhibition of GSDMD-capase 8 pathway. These results might provide important novel insights into the biological roles of IRE-1α in hepatic pathophysiology and hence provide new therapeutic targets for drug discovery in the future.

## Materials and Methods

### Animals

C57BL/6J SPF male mice between 6 and 8 weeks old were purchased from DASHUO Animal Company (SCXK (CHUAN) 2015–30, Chengdu, China). Mice were gavaged with chow (DMSO) with or without intraperitoneal injection of STF083010 (30 mg/kg body weight) only once purchased from MedChemExpress (MCE, CN). Body weight and temperature were monitored every day. All of the experimental procedures were approved by the Institutional Animal Care and Use Committee (IACUC) of Sichuan University. The detailed information of animal models was described in the corresponding parts of results section.

### Cell culture

HUVECs were cultured using the EndoGRO-VEGF Complete Culture Media (ECM) Kit (Millipore, Billerica, MA, USA). The human hepatocyte line L-02, colon cancer HT-29, lung cancer A549, kidney HK-2, mouse hepatocyte AML-12, myoblast C2C12, intestinal epithelial IRD98, and cardiomyocyte HL-1 cells (purchased from ATCC) were cultured in high glucose DMEM supplemented with 10% fetal bovine serum (FBS) and 1% antibiotics (100 U/ml penicillin and 100 μg/ml streptomycin) at 37°C. When reaching approximately 80–90% confluence, the cells were digested and cultured overnight in 96-well plates or 6-well plates. Then, the medium was changed to fresh medium containing 10% FBS and different treatments. 4u8c and APY29 are specific IRE-1α inhibitors purchased from MedChemExpress (MCE, CN). Lipopolysaccharide (LPS) and ATP were purchased from Topscience Biotechnology (Topscience, Shanghai, CN). Tunicamycin (TM) and thapsigargin (Thp) are chemical ER stressors purchased from Santa Cruz Biotechnology (Santa Cruz, CA, USA). After treatments, the medium was collected, and the cells were harvested for subsequent assays.

### Blood biochemical analysis

Serum alanine transaminase (ALT), aspartate aminotransferase (AST), urea (UREA), uric acid (UA), creatinine (CREA), blood urea nitrogen (BUN) and total bilirubin (TB) were detected by an autoanalyzer (Cobas 6000 c501, Roche Diagnostics, Switzerland).

### Histological analysis

All samples were sliced into 5 μm sections and were routinely stained with hematoxylin and eosin (H&E). The 10 μm frozen liver sections were stained with Oil Red O (Nanjing Jiancheng Bioengineering Institute, Nanjing, China) to visualize lipid droplets in the liver. TUNEL staining of the tissue sections was conducted using the DeadEnd Fluorometric TUNEL System (Promega), followed by immunofluorescence staining with DAPI. The histological features were captured by a Vectra Polaris™ Automated Quantitative Pathology Imaging System (PerkinElmer, USA).

### RNA isolation and quantitative real-time PCR

Total RNA was isolated from cultured cells or fresh liver samples using TRIzol reagent (Ambion, USA), and 1.0 μg RNA was reverse-transcribed into cDNA using a High-Capacity cDNA Synthesis Kit (Vazyme, China) according to the manufacturer’s protocol. Real-time PCR was performed to assess gene expression in a Bio-Rad QPCR Machine using SYBR Green master mix (Vazyme, China). Each sample was amplified in triplicate, and the expression of β-actin was used as an internal control for every PCR assay. The primer sequences used for qRT−PCR are shown in Supplementary Table S1.

### Western blot analysis

Total protein was extracted from cultured cells or fresh liver samples using RIPA lysis buffer. A BCA Protein Assay Kit (Cwbio, China) was used to measure protein concentrations. Protein extracts (60 μg) were separated on a 10% SDS–PAGE gel and transferred to a 0.2 μm PVDF membrane. The immunoblots were visualized using a ChemiDoc™ imaging system (BioRad, USA). The antibodies used for western blot are shown in Supplementary Table S2.

### Enzyme-linked immunosorbent assay (ELISA)

Cell culture supernatants or serum were measured for IL-1β (ab46052; Abcam,USA), TNF-α (A098526-48T; Affandi, CN), and K^+^ (BC2775; Solarbio, CN) using ELISA kits according to the manufacturer’s instructions.

### Immunofluorescence analyses

Immunofluorescence staining was performed to detect the expression of target proteins via specific antibodies. DAPI was used to stain the nucleus of cells. Images were visualized on an SP5 confocal microscope (Leica Microsystems) using excitation lasers and processed using ImageJ (National Institutes of Health).

### Plasmids and siRNAs

JetPRIME® (Polyplus) was used as the transfection reagent for the initial set of plasmids and siRNA experiments (according to the manufacturer’s instructions). Plasmids were purchased from Vigene Biosciences (MD, USA). All siRNAs were purchased from Hanheng Biotechnology (Shanghai, CN). Sequences of siRNAs used in this study were as follows: SiContril sense strand, 5’-UUCUCCGAACGUGUCACGUTT-3’; antisense strand, 5’-ACGUGACACGUUCGGAGAATT-3’; siMDMX sensestrand, 5’ CCAUUAUCCUGAGCACCUUTT-3’; antisense strand, 5’-AAGGUGCUCAGGAUAAUGGTT-3’.

The cells were plated prior to transfection such that they were only 80% confluent prior to RNA isolation. RNA was extracted 24 h after cell transfection.

### Acridine orange and ethidium bromide (AO/EB) staining

The cells were treated with BMSO, STF083010, overexpression plasmid and ATP plus LPS for an additional 24 h and then subjected to AO/EB staining (100 μg/ml AO and 100 μg/ml EB mixed in PBS, Aladdin, China). The fluorescence of the cells was observed under a fluorescence microscope.

### Statistical analysis

Experiments were performed at least three times, and quantitative data are expressed as the mean ± SD. GraphPad Prism 6 was used for statistical analyses. Data were evaluated with a 2-tailed, unpaired Student’s t test or compared by one-way analysis of variance. A value of P < 0.05, **P < 0.01 and ***P < 0.001 was considered statistically significant.

## Acknowledgement

This study was supported by the Program of National Natural Science Foundation of China (Chengdu, China, 82170590, 81870609, and 82170075), Science and Technology Department of Sichuan Province project funding (2021YFH0061), and the 1.3.5 project for disciplines of excellence, West China Hospital, Sichuan University (Chengdu, China, ZYGD18014).

## Supplemental material

**Figure 1.**
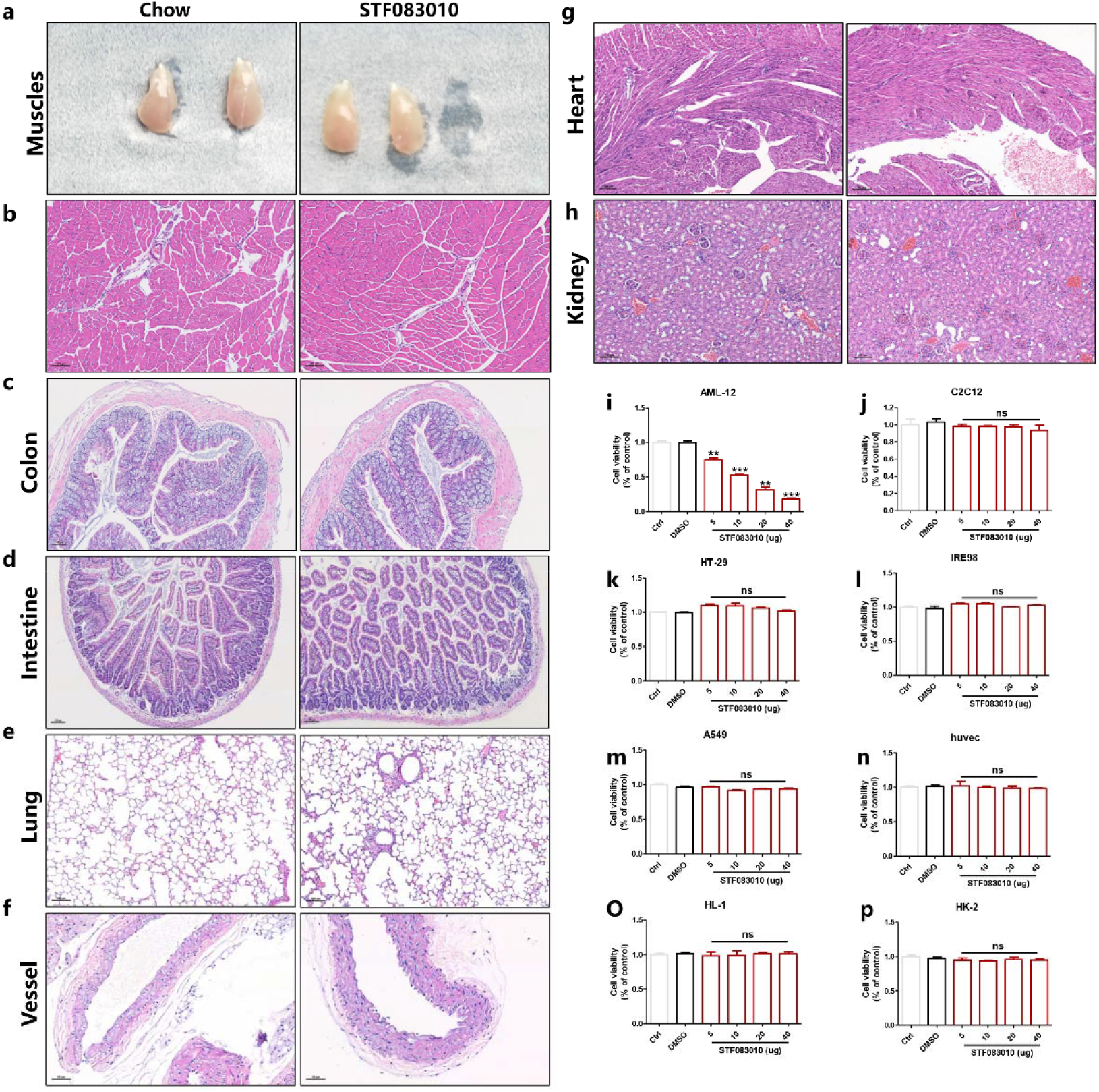
Inhibition of IRE-1α not lead to other tissues injury. a Morphology of skeletal muscle in mice b-h Representative H&E-staining. i-p Viability of cells was treated with STF083010 for 24 hours. Data information: Data are presented as means ± SD. **P < 0.01, determined by ANOVA.

**Figure 2.**
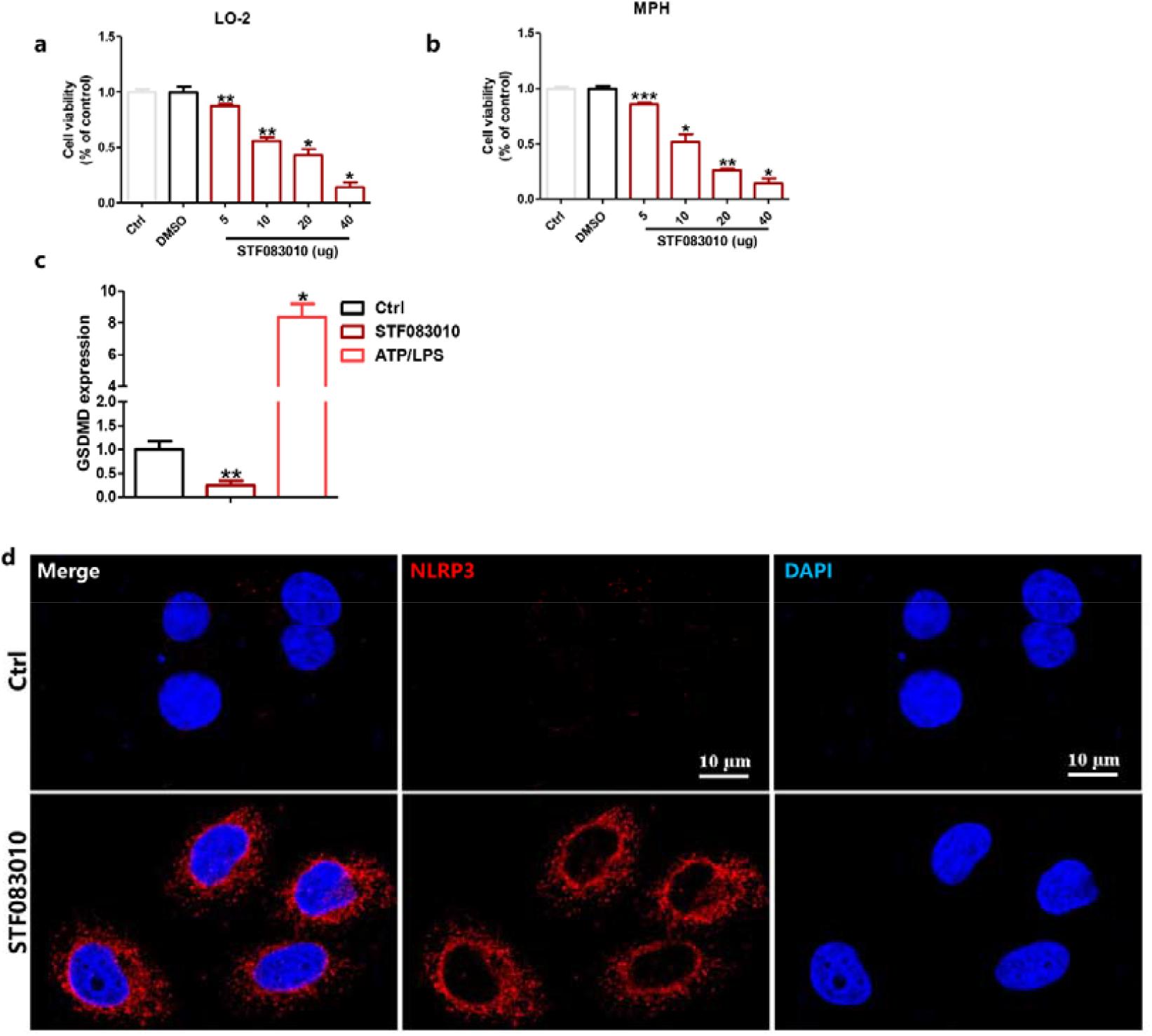
GSDMD is a negative executor of necroptosis but NLRP3. a, b Viability of cells was treated with STF083010 for 24 hours. c the mRNA expression of STF083010–treated LO-2 24 h. d Cells were exposed to STF083010 and subsequently labeled by immunofluorescence with anti-NLRP3 (red) and DAPI (blue) (scale bars; 10 μm), respectively. Data information: Data are presented as means ± SD. *P < 0.05, **P < 0.01, ***P < 0.001, determined by ANOVA.

**Figure 5.**
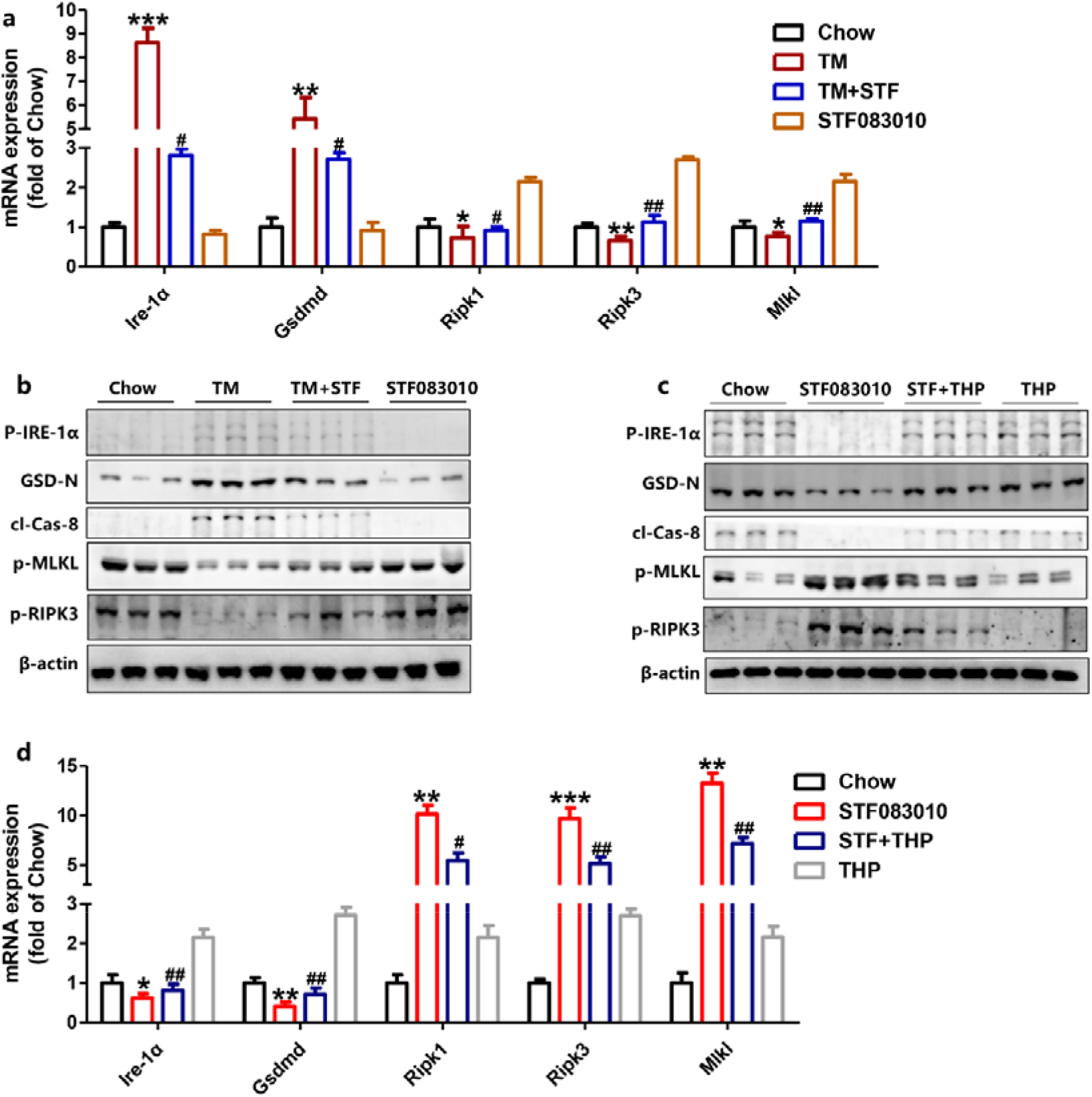
IRE-1α equilibrium improves mice survival. a, d the mRNA expression of Ire-1α, Gsdmd, Ripk1, Ripk3 and Mlkl was detected after different treatment, and β-actin was used as an internal control. b, c Representative western blots of key proteins after different treatment, and β-actin was used as a protein-loading control.

**Table S1.**
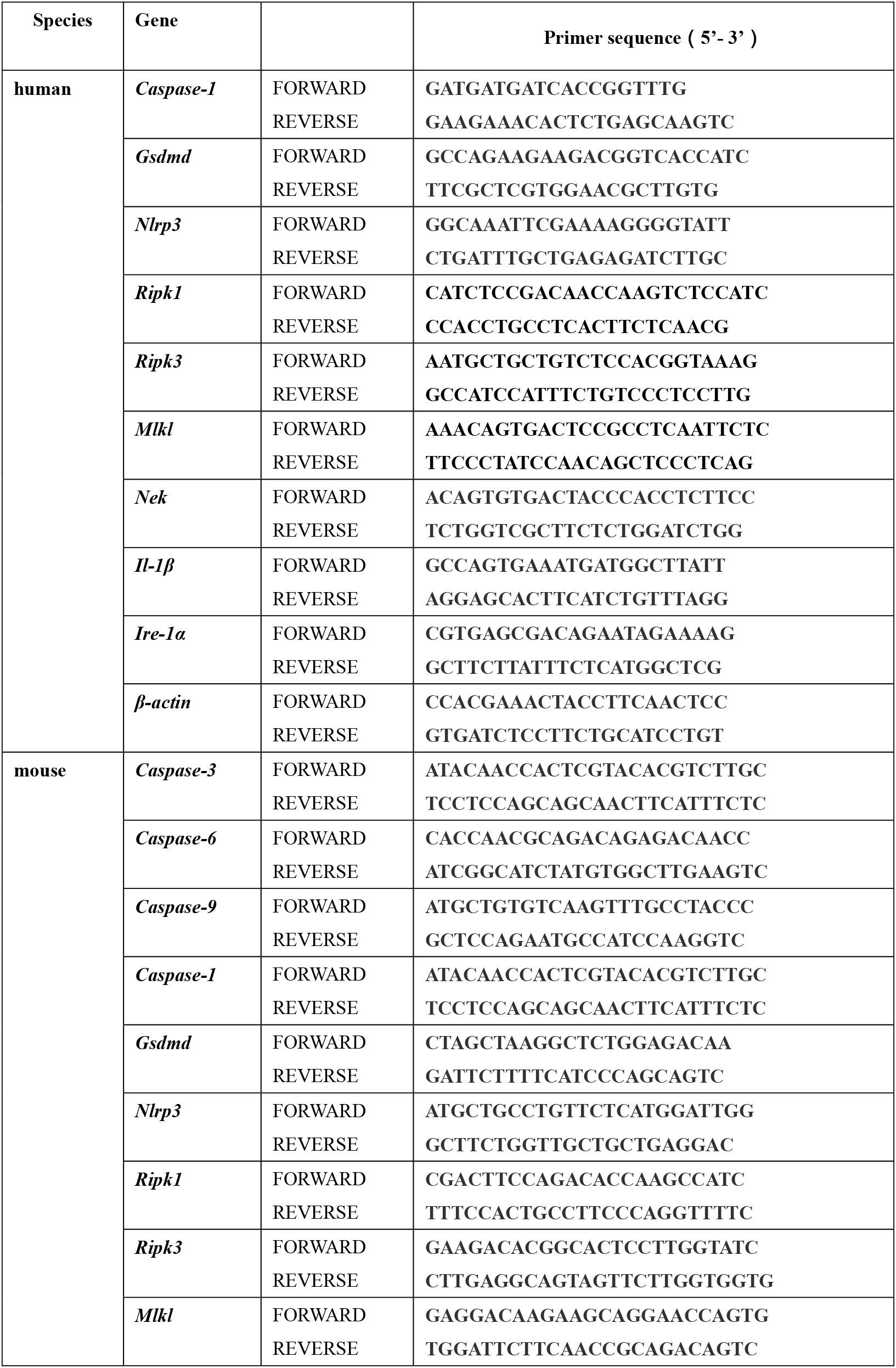
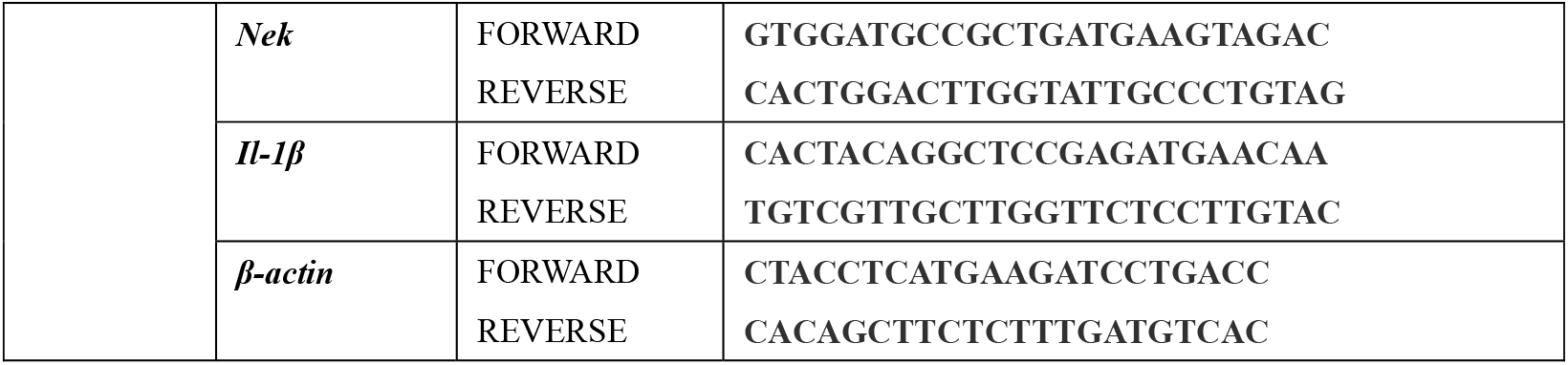
Primer sequences for real time-PCR.

**Table S2.** Antibodies for Western Blot and Immunofluorescence.

| **Antibodies** | **Source** | **Identifier** |
| --- | --- | --- |
| Mouse monoclonal HRP-conjugated Beta actin | Proteintech Group | Cat#: HRP-60008; |
| Cleaved Caspase-3 (Asp175) (5A1E) Rabbit | Cell Signaling Technology | Cat#: #9664 |
| Cleaved Caspase-6 (Asp162) Rabbit | Cell Signaling Technology | Cat#: #9761 |
| Cleaved Caspase-9 (Asp353) Rabbit | Cell Signaling Technology | Cat#: #9509 |
| Cleaved Caspase-1 (Asp297) (D57A2) Rabbit | Cell Signaling Technology | Cat#: #4199 |
| NLRP3 (D4D8T) Rabbit | Cell Signaling Technology | Cat#: #15101 |
| Cleaved Gasdermin D (Asp275) (E7H9G) Rabbit | Cell Signaling Technology | Cat#: #36425 |
| Caspase-8 (D35G2) Rabbit | Cell Signaling Technology | Cat#: #4790 |
| Phospho-MLKL (Ser345) (D6E3G) Rabbit | Cell Signaling Technology | Cat#: #37333 |
| Phospho-RIP3 (Ser227) (D6W2T) Rabbit | Cell Signaling Technology | Cat#: #93654 |
| Phospho-IRE-1(Ser724) Rabbit | Invitrogen | Cat#: PA5-105424 |

